# The gut virome of healthy children during the first year of life is diverse and dynamic

**DOI:** 10.1101/2020.10.07.329565

**Authors:** Blanca Taboada, Patricia Morán, Angélica Serrano-Vázquez, Pavel Iša, Liliana Rojas-Velázquez, Horacio Pérez-Juárez, Susana López, Javier Torres, Cecilia Ximenez, Carlos F. Arias

## Abstract

In this work, we determined the diversity and dynamics of the gut virome of infants during the first year of life. Fecal samples were collected monthly, from birth to one year of age, from three healthy children living in a semi-rural village in Mexico. Most of the viral reads were classified into six families of bacteriophages including five dsDNA virus families of the order *Caudovirales*, with *Siphoviridae* and *Podoviridae* being the most abundant. Eukaryotic viruses were detected as early as two weeks after birth and remained present all along the first year of life. Thirty-four different eukaryotic virus families were found, where eight of these families accounted for 98% of all eukaryotic viral reads: *Anelloviridae*, *Astroviridae*, *Caliciviridae*, *Genomoviridae*, *Parvoviridae, Picornaviridae*, *Reoviridae* and the plant-infecting viruses of the *Virgaviridae* family. Some viruses in these families are known human pathogens, and it is surprising that they were found during the first year of life in infants without gastrointestinal symptoms. The eukaryotic virus species richness found in this work was higher than that observed in previous studies; on average between 7 and 24 virus species were identified per sample. The richness and abundance of the eukaryotic virome significantly increased during the second semester of life, probably because of an increased environmental exposure of infants with age. Our findings suggest an early and permanent contact of infants with a diverse array of bacteriophages and eukaryotic viruses, whose composition changes over time. The bacteriophages and eukaryotic viruses found in these children could represent a metastable virome, whose potential influence on the development of the infant’s immune system or on the health of the infants later in life, remains to be investigated.

## Introduction

Humans are in constant close contact with a rich variety of bacteria, archaea, fungi, protists and viruses, which all together reside on, or within the human body, forming an ecosystem denominated microbiota. Distinct body sites are characterized by distinct microbial communities, with the gastrointestinal tract being the most populated site, playing an important role in human health and development. The bacterial component of the microbiota is the best characterized, having many functions in host physiology like metabolic processes, development and maturation of the immune system [1,2]. Bacterial colonization begins during birth and continues to change and evolve throughout life and its composition can be influenced by factors as birth mode, gestational age, antibiotic usage, diet, geographical location, lifestyle and age [3,4]. Modifications in the microbiota composition can lead to several diseases, including obesity, diabetes, or cardiovascular diseases, among others

Contrary to bacterial composition, the virus assembly (known as virome) in the gut is much less understood. It has been observed that bacteriophages are the predominant viruses, where *Siphoviridae*, *Inoviridae*, *Myoviridae*, *Podoviridae* and *Microviridae* represent the most common families [5–7]. Eukaryotic viruses have also been commonly found in the gut of healthy individuals [8], with animal viruses of the *Anelloviridae*, *Picobirnaviridae*, and *Circoviridae* families being the most frequently reported [8], although viruses from the *Adenoviridae*, *Astroviridae*, *Caliciviridae*, *Parvoviridae*, *Picornaviridae*, and *Polyomaviridae* families have also been described [5–7,9]. In addition, the presence of different families of plant viruses, including A*lphaflexiviridae*, *Tombusviridae*, *Nanoviridae*, *Virgavirida*e and *Geminiviridae*, have also been commonly reported.

In this work, the viral composition of the gastrointestinal tract of three healthy infants from a semi-rural town in Mexico was characterized in fecal samples collected monthly during their first year of life. The presence of eukaryotic viruses was detected as early as two weeks after birth and represented a diverse metastable and dynamic virome along the year of study formed by a rich and abundant mixture of viruses from plant and animal hosts.

## Materials and methods

### Population studied and sample collection

This study was carried out in the semi-rural community of Xoxocotla, in the state of Morelos, 130 km south of Mexico City. Healthy women who arrived for routine control at the pregnancy clinic of the Xoxocotla Health Center, during the last trimester of pregnancy, were invited to participate in the study. Written informed consent was obtained from all mothers after providing them with detailed information about the study and its characteristics. The protocol and the consent letter were approved by the Scientific and Ethics Committee of the Medical School of the National University of Mexico as well as by the Ministry of Health of the State of Morelos. Three mother-infant pairs were included in this study; the infants were healthy, full-term products, without any congenital condition and with normal weight at birth. A single fecal sample was obtained at the end of the last trimester of pregnancy from each mother. The infants were followed monthly during their first year of life between March 2015 and June 2016; all stool samples were taken under aseptic conditions to avoid contamination. The samples were not exposed to antiseptics or disinfectants. All samples were collected from the diaper in sterile containers, transported to the lab in cold chain and frozen at −70 °C until use.

### Nucleic acid isolation and sequencing

Nucleic acids were extracted from the stool samples as described before [10]. Briefly, 10% stool homogenates were prepared in phosphate-buffered saline (PBS); the, chloroform (10%) and 100 mg of 150 to 212 μm glass beads (Sigma, USA) were added in final volume of 1 ml and processed in a bead beater (Biospec Products, USA). The samples were centrifuged at 2000 x g to remove large debris, and the recovered supernatants filtered through Spin-X 0.45 μm pore filters (Costar, NY). A volume of 400 μl of filtered samples was treated with Turbo DNAse (4 U) (Ambion, USA) and RNAse (0.3 U) (Sigma, USA) for 30 min at 37°C. Nucleic acids were extracted using the PureLink viral RNA/DNA extraction kit according to the manufacturer’s instructions (Invitrogen, USA), and eluted in nuclease-free water, aliquoted, and stored at −70°C until further use. Nucleic acids were random amplified with SuperScript III reverse transcriptase (Invitrogen, USA) with primer A (5’– GTTTCCCAGTAGGTCTCN_9_-3’). The cDNA was generated by two consecutive rounds of synthesis with Sequenase 2.0 (USB, USA). The synthesized cDNA was then amplified with Phusion High fidelity polymerase (Finnzymes) using primer B (5’-GTTTCCCAGTAGGTCTC-3’) and 10 additional cycles of the program: 30 sec at 94°C, 1 min at 50°C and 1 min at 72°C. Then, the DNA was purified using ZYMO DNA Clean & Concentration-5 kit. Sequencing libraries were prepared using Nextera XT DNA library preparation kit (Illumina); samples were uniquely tagged, pooled and then deep sequenced on the Illumina NextSeq500 system, generating paired-end reads of 75 bases. The base calling was performed by Illumina Real Time Analysis (RTA) v1.18.54 software and the demultiplexing of reads by bcl2fastq v2.15.0.4.

### Metagenomic data analysis

A viral metagenomics pipeline, which includes quality controls and filtering, taxonomic annotation was applied as previously described [11]. Briefly, the process was: i) Quality control, adapters and low quality bases from 5’ and 3’ ends were trimmed, low complexity reads or shorter than 40 bases were removed using Cutadapt v1.16 [12] and exact duplicates reads were excluded by using CD-HIT-DUP v.4.6.8 [13]. Finally, ribosomal RNA and human genome reads were filtered by aligning them against ribosomal sequences from Silva database (DB) [14] and human genomes sequences from GenBank, using SMALT v0.7.5 [15]. The remaining sequences were considered valid reads. ii) Taxonomic classification, valid reads were mapped to a viral reference DB obtained from nt (minimally non-redundant nucleotide database) from NCBI [16], using SMALT at 70% of identity, and mapped reads were assembled using IDBA-UD software [17]. Not assembled contigs and singlets reads were compared against nt database using BLASTn [18] to remove false positives. The same process was done to identify bacteria and fungi. Then, the reads that did not map using the nucleotide alignment were assembled using IDBA-UD and contigs greater than 200 bases were compared to all proteins of nr (minimally non-redundant protein database) using BLASTx. Then, the software MEGAN 5.10.6 [19] was used to assign reads and contigs to the most appropriate taxonomic level.

### Statistical analysis

Nucleotide and protein taxonomic reads assigned to viruses were extracted from MEGAN to generate a count matrix. Differences in the sequencing depth of the samples were corrected by dividing the number of bacteria and viruses reads by each sample valid reads and normalized to 5 million. In addition, the taxon abundance differences were analyzed with the Trimmed Mean of M-value (TMM) method [22]. Unless otherwise indicated, statistical analyses were conducted in R-3.5.3 statistical environment [23], using the Vegan package [24]. To assess the alpha diversity, we calculated richness as the expected number of species, Shannon diversity (H) index and Pielou’s on count matrix. For beta diversity, Bray–Curtis distance metric was used [25]. The distances were used as input for the Nonmetric Multidimensional Scaling (NMDS) ordination method. For comparison of groups, samples were divided into cases and controls and in order to associate them with metadata factors, a nonparametric multivariate permutation test (PERMANOVA) analysis was done using the Adonis function with 999 permutations, and Mann-Whitney’ test for measures [26]; homogeneity variances between groups were verified in all comparisons. Finally, the differential abundance of taxons was carried out with the EdgeR package (version 3.24.3) in R (version 3.5.3), as described in Loraine et al. [27]. Common and tag-wise methods were used to estimate the biological coefficient of variation and “exact” test used to perform hypothesis testing and false-discovery rates to adjusted p-values. All statistics were considered significant if p < 0.05.

### Data availability

We submitted to the NCBI database the Bioproject PRJNA592261 which contains all the biosamples and complete genomes.

## Results

### Infant’s cohort and sample analysis

Fecal samples from three apparently healthy infants, with no disease symptoms during the study, were collected monthly, starting two weeks after birth and until 12 months of age, obtaining 11 samples from infant 2, and 12 samples from infants 4 and 5. Mother samples were taken at a single time point around the eighth month of pregnancy. The characteristics of mothers and infants are described in S1 Table. Two infants were born via cesarean section (infant 4, male; infant 5, female), while infant 2 (male) was born via vaginal delivery. The three infants were breastfed all along the year of study, although they received supplemental formula after their first three months of life. Only the mother of infant 2 was exposed to antibiotics before sample collection, but none of the infants received antibiotics throughout the study.

Total nucleic acids were extracted from 38 fecal samples (35 from infants and 3 from mothers) to detect both DNA and RNA viruses and sequenced using the paired-end 2×75 bp Illumina HiSeq 500 system (Illumina, Inc). Initially, 421.78 million paired-end reads were obtained for all libraries, with a mean of 12.05 ± 3.23 millions per sample (S2 Table). After quality control, duplication removal and filtering (human and ribosomal) processes, 90.84 million valid reads remained (mean= 2.39 millions, ranging from 0.68 to 3.58).

### Viral taxonomy composition

On average, it was found that 36.1% (±27.6% s.d.) and 26.9% (±14.3% s.d.) of the valid reads in the infant samples had homology to at least one viral or bacterial reference sequence, respectively (Fig 1). In contrast, in mothers only 4.2% (±2.4% s.d.) of valid reads were identified as viral, whereas bacterial reads were as high as 47.9% (±5.0% s.d.). Similar to previous viral metagenomics studies, 37.8% (±17.8% s.d.) of valid reads per sample showed no significant similarity to any known sequence of GenBank DB [28,29]. Reads mean percentages were calculated for each sample first and then, the global mean for all samples was estimated.

**Fig 1.**
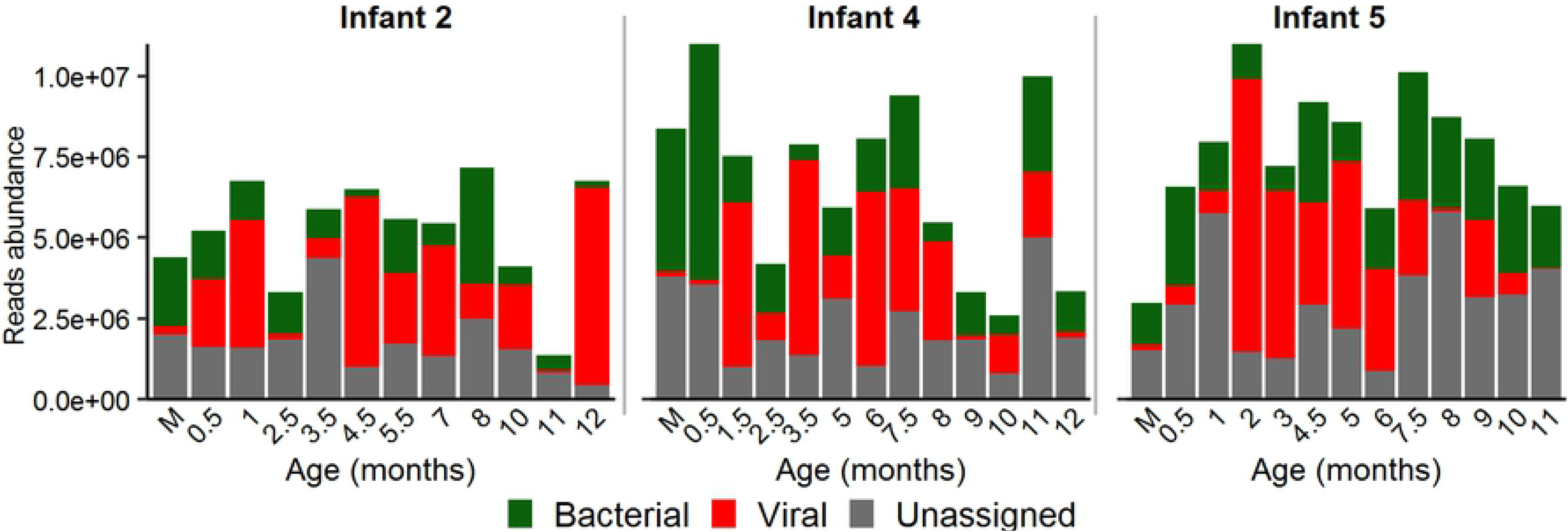
Abundance of bacterial and viral sequence reads in each sample. Samples from infants are indicated according to their age in months and from mothers as M.

### Eukaryotic viruses

In infants, 15.5% of the assigned viral reads correspond to eukaryotic viruses, which can be classified within 33 families (Fig 2). These include 15 virus families with single-stranded RNA genomes (ssRNA), 4 with double-stranded RNA genomes (dsRNA), 1 with ssRNA (reverse transcriptase) genome, 5 with ssDNA genomes and 8 with dsDNA genomes. Regarding the hosts of these virus families, about half of them infect vertebrates (42.4%), followed by those that infect plants (21.2%), plants/fungi (12.2%), invertebrates (12.2%), amoebae (6.0%), algae (3.0%) and fungi (3.0%) (Fig 2). It is important to point out that nine of the viral families identified accounted for 97% of all reads: i) *Virgaviridae*, the most abundant, was identified in all samples (35/35), representing an average of 26.6% of all eukaryotic viral reads and ranging from 0.01% to up to 99.7% of all reads in the various samples. ii) *Anelloviridae* and *Picornaviridae* represented on average 24.8% and 18.7% of all eukaryotic viral reads, respectively, and were found in up to 80% of the samples (28/35). iii) *Caliciviridae*, *Parvoviridae* and *Reoviridae* were less abundant, on average 8.8%, 5.8% and 5.8%, respectively, and less prevalent, been found in 66% (23/35), 71% (25/35) and 60% (21/35) of samples, respectively. iv) *Astroviridae* with 3.3% of abundance, and *Genomoviridae* with 0.5%, were identified in 43% (15/35) and 49% (17/35) of the samples, respectively. v) *Circoviridae* had around 3.3% of abundance and was identified in only 17% (6/35) of the samples. As expected, the most abundant and prevalent viral families found have humans as hosts, except for the family *Virgaviridae*, whose natural hosts are plants. On the other hand, among the viral families identified in a single sample or in less than 10% of the samples, only 15% infect invertebrates. In the samples from the three mothers, 18.6% of the viral reads were eukaryotic, classified into 16 families, and the rest were bacteriophages, with *Virgaviridae* (90.6%) and *Phycodnaviridae* (8.3%) as the dominant virus families.

**Fig 2.**
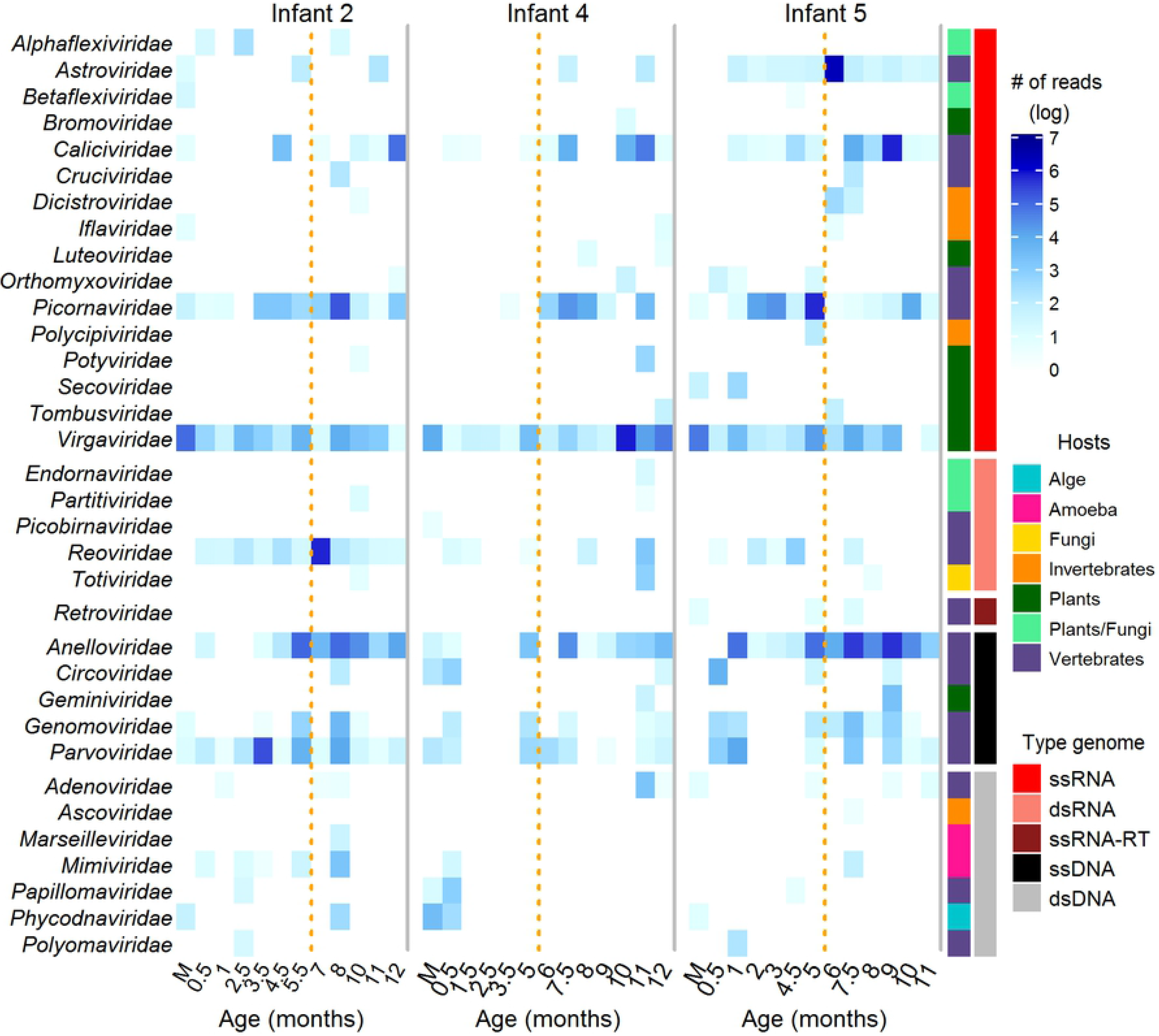
Read distribution of eukaryotic viral families in infants during the first year of life, and their mothers. Samples from infants are indicated according to their age in months and the single mother sample in each case, as M. The virus family name is indicated on the left side, while the host and type of virus genome are shown in the right side. The discontinuous yellow lines divide the years into two semesters.

At the genus level, 54 viral genera were identified in the infant samples and 18 were present in the samples obtained from their mothers (Fig 3). *Tobamovirus* of the *Virgaviridae* family was the most prevalent genus, being found in all mother and infant samples, and was also the most abundant in infants, representing in average 28.5% of the eukaryotic viral reads per sample. It was also the predominant viral genus in mothers, representing in average up to 91.0% of all virus reads. A detailed characterization of the plant viruses found in the infants’ gut was reported previously by our group [30]. From the remaining eukaryotic viruses, *Betatorquetenovirus* (*Anelloviridae*), *Bocaparvovirus* (*Parvoviridae*), *Enterovirus* (*Picornaviridae*), *Norovirus* (*Caliciviridae*) and *Rotavirus* (*Reoviridae*) were also abundant genera representing, in average, 15.5%, 6.2%, 11.5%, 7.8% and 6.2% of the reads in the infant samples, respectively, and were detected in more than 65% of the fecal samples. *Parechovirus* (*Picornaviridae*) and *Mamastrovirus* (*Astroviridae*) were less abundant, representing 6.9% and 3.1% of the viral reads, respectively and found in only 40% of the samples. While *Cardiovirus* (*Picornaviridae*), *Cyclovirus* (*Parvoviridae*) and *Sapovirus* (*Caliciviridae*) were identified only sporadically. Of note, more than half of the genera were uniquely identified in a single sample of some of the infants (Fig 3.).

**Fig 3.**
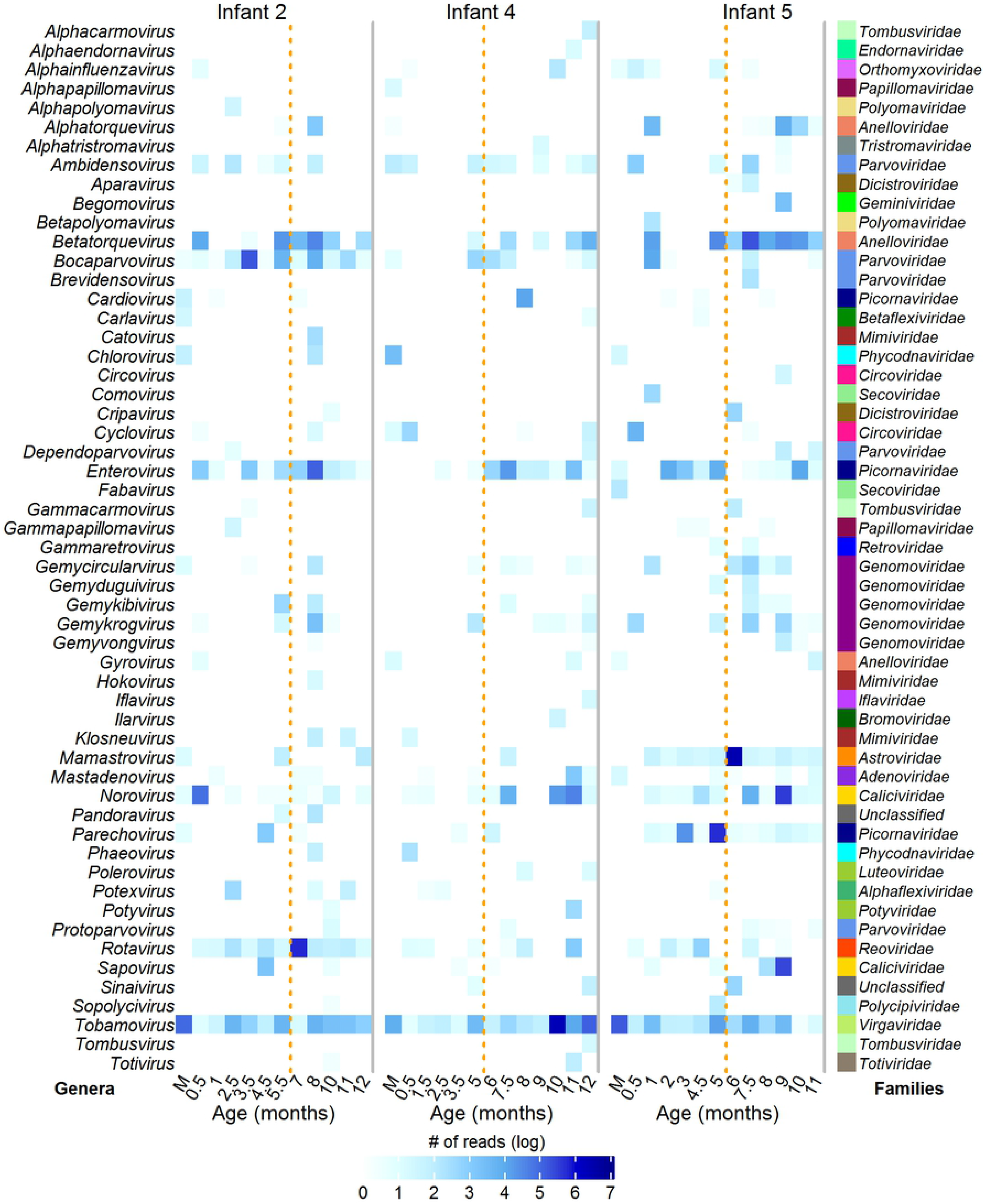
Distribution of eukaryotic viral genera reads in infants during the first year of life and their mother. Samples from infants are indicated according to their age in months and, from mothers, as M. The name of the viral genera is indicated on the left side while the family names are shown on the right side. The discontinuous yellow lines divide the year into semesters.

In contrast to previous studies, we were able to classify reads at lower taxonomic levels, identifying 124 eukaryotic viral species in infants and 30 in mothers (S3 Table). On average, 15 (±9 s.d.) viral species were identified per infant sample and 12 (±1) in each mother’s sample. Most species were identified sporadically, with 70% of them being observed in only 10% of infant’s samples. However, some species were frequently found in samples throughout the year. Fig 4 shows the most common and abundant species across all samples. Interestingly, tropical soda apple mosaic virus (TSAMV) and pepper mild mottle virus (PMMoV), from the plant infecting *Virgaviridae* family, were the most prevalent and abundant species in both, infant and mother samples. TSAMV was found in all samples, representing on average 15.9% of the eukaryotic viral reads per infant sample (range 0.001-89.4%) and 62.9% of reads in the mothers’ samples (range 0.7-95.9%). Whereas PMMoV represented, in average, 10.3% of the viral reads in the infant samples and 3.5% in the mothers’ samples; this virus was detected in 80% and 100% of infant and mother samples, respectively. Other abundant virus species found in more than half of the infant’s samples, were Norwalk virus (NV), rotavirus A (RVA), and torque teno virus (TTV), with 6.9%, 6.1 % and 6.0% of abundance, respectively. Other prevalent viral species identified in more than 30% of the samples were chimpanzee torque teno mini virus (Cpz TTMV), human astrovirus 1 (HAstV 1), enterovirus A (CVA-2), rhinovirus A (RV-A1), parechovirus A (HPeV), rattail cactus necrosis-associated virus (RCNaV), torque teno midi virus (TTMDV) and TTV-like mini virus (TLMV), ranging in abundance from 1.1% to 6.2% per sample. Finally, enterovirus B (CVB 3) and Sapporo virus (SV) species were present in 15% of the samples with an abundance of 2%.

**Fig 4.**
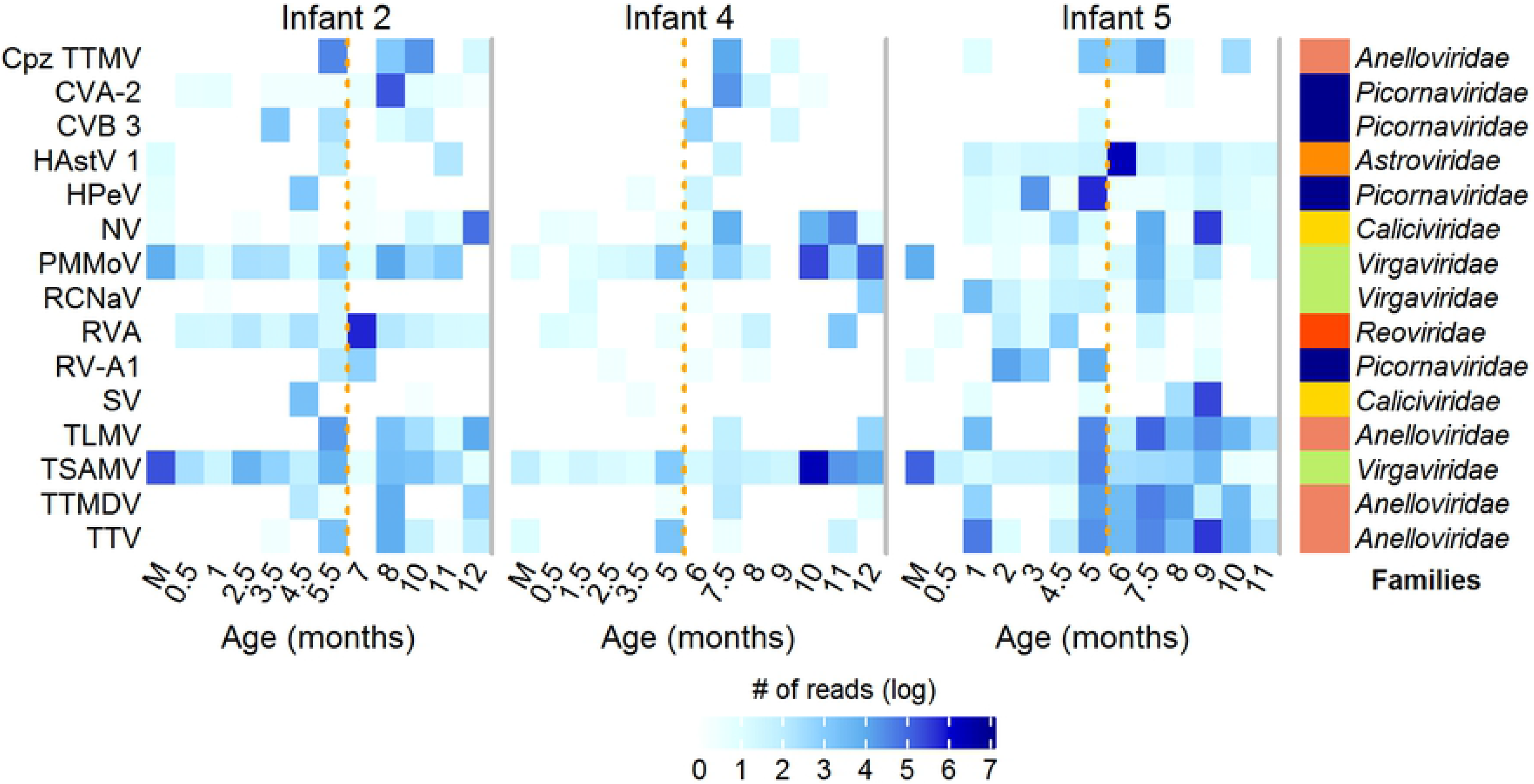
Most abundant and prevalent eukaryotic viral species in infants during the first year of life and their mothers. Samples from infants are indicated according to their age in months and from mothers as M. The family to which each virus species belongs is indicated on the right side. *Cpz TTMV, chimpanzee torque teno mini virus; CVA-2, enterovirus A; CVB 3, enterovirus B; HAstV 1, human astrovirus 1; HPeV, parechovirus A; NV, Norwalk virus; PMMoV, pepper mild mottle virus; RCNaV, rattail cactus necrosis-associated virus; RVA, rotavirus A; RV-A1, rhinovirus A; SV, Sapporo virus; TLMV, TTV-like mini virus; TSAMV, tropical soda apple mosaic virus; TTV, torque teno virus and TTMDV, torque teno midi virus. The discontinuous yellow lines divide the year into semesters.

### Bacteriophages

As previously observed [5,7,29,31–33], the vast majority of viral reads identified were classified into six different families of bacteriophages, with 84.5% of abundance in infants and 81.4% in mothers (Fig 5). In infants, five dsDNA families of the order *Caudovirales* were the most abundant: *Siphoviridae* (long, non-contractile tailed-phages, temperate, with some lytic members), with an average of read abundance of 51.5% per sample; *Podoviridae* (short, contractile tailed-phages, lytic, with some temperate members) with 20.2%; *Myoviridae* (long, contractile tailed-phages, strictly lytic) with 12.7%; the provisional family crAss-like (contractile tailed-phages, strictly lytic, with podovirus-like morphology) with 10.4%; and *Ackermannviridae*, the new bacteriophage family proposed in 2017, with 2.9%. The ssDNA family *Microviridae* (lytic, with some identified as prophages) represented, on average, 2.3% of children’s bacteriophage reads. Interestingly, in mothers, the crAss-like bacteriophages predominated in the gut, with an average of 79% of the total phage reads.

**Fig 5.**
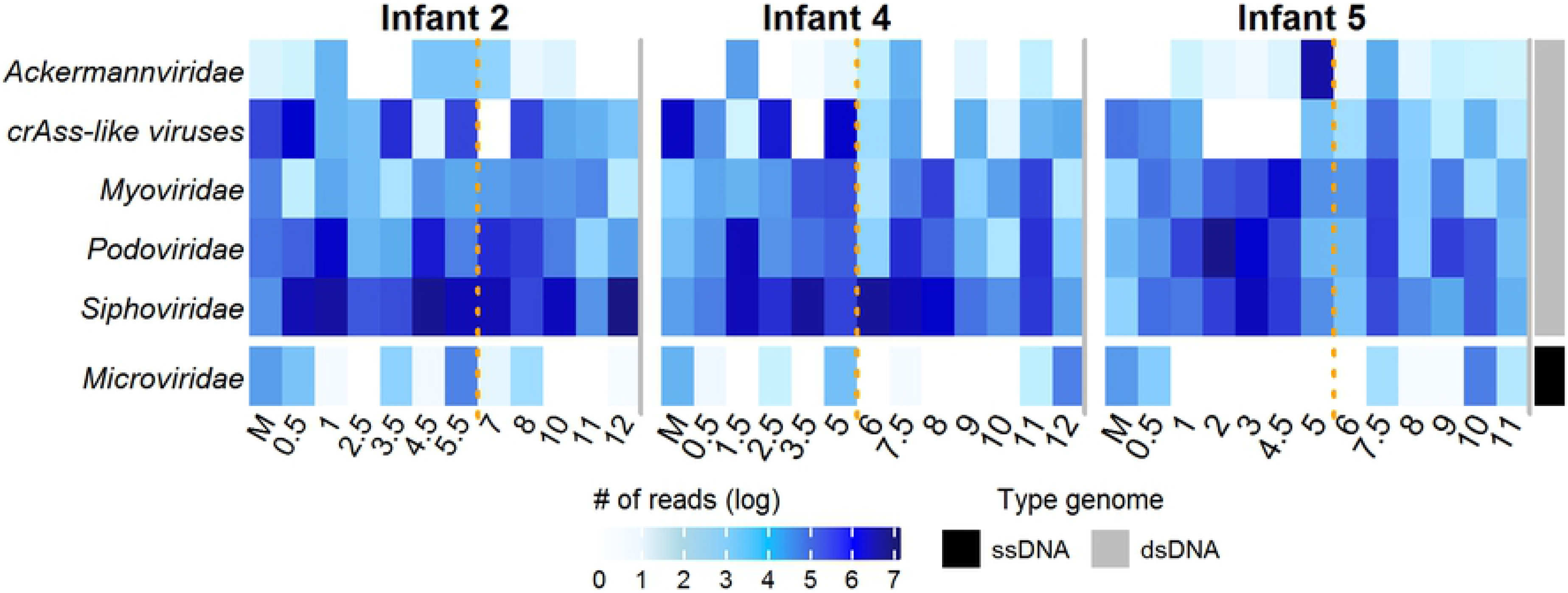
Abundance of families of bacteriophages in infants during the first year of life and their mothers. Samples from infants are indicated according to their age in months and from mothers as M. The type of genome of each viral family is indicated on the right side. The discontinuous yellow lines divide the year into semesters.

Regarding phage genera, 97 were identified in infants and 31 in mothers (S4 Table), with the genus *Pis4avirus* from the *Siphoviridae* family and the genus *G7cvirus* from the *Podoviridae* family being the most common, since they were identified in all infant samples. Other frequent and abundant genera identified in more than 75% of samples were: *Sk1virus*, *Jerseyvirus*, *K1gvirus* and *Lambdavirus* of the *Siphoviridae* family; *T7virus*, *P22virus* and *Epsilon15virus* of the *Podoviridae* family and *P1virus* of the *Myoviridae* family. On the other hand, more than 31 genera were identified in a single sample or in less than 3 in the infant’s samples.

In contrast with previous reports that have shown that a stable community of bacteriophages exist in adults over a long period of time [34,35], we observed a dynamic and unstable community of phages in infants during their first year of life (S5 Table), in agreement with previous studies in early childhood [5,36]. The majority were identified in a single sample with 48% of the phages (266 species) found in only one or two samples. However, we found 82 species shared in one third of the infant samples, 29 species in 50%, and 7 species in more than 80% of the samples. This more stable phageome included phages of high abundance, e.g., the 7 most frequently species also accounted for 33% of the total read abundance (Fig 6). On average, infants harbored 79 species per sample (range 36-194), 18 from the *Myoviridae* and *Podoviridae families*, 40 from the *Siphoviridae* and 1 from the *Ackermannviridae*, crAss-like and *Microvirid*.

**Fig 6.**
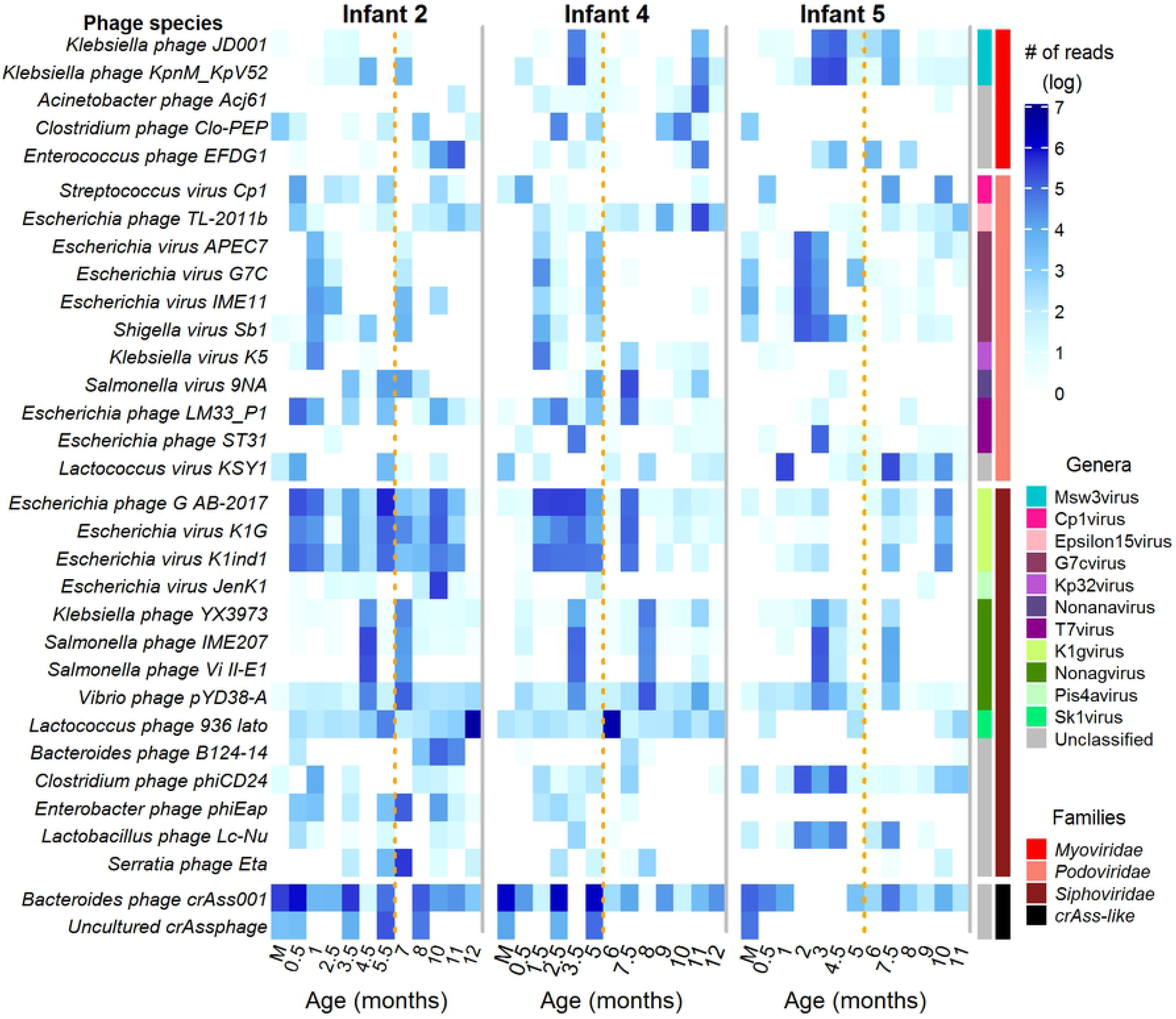
Most abundant and prevalent phage species in infants during the first year of life and their mothers. Samples from infants are indicated according to their age in months and from mothers as M. The family for each phage species is indicated on the right side. The name of the phage species is listed on the left side. The discontinuous yellow lines separate the semesters.

The most abundant and frequent phage species in both infants and mothers has around 75% of homology to the Bacteroides phage crAss001 from the crAss(cross Assembly)-like family (Fig 6), which has been recently reported to be the most abundant and prevalent type of crAssphage in the adult human gut, and to infect bacteria of the order *Bacteroidales* [37]. This phage species was found in all mothers’ samples and in 87% of the infants’ samples, with an average abundance of 71.7% and 13.7% of total phage reads per sample, in mothers and infants, respectively. Other common and abundant bacteriophages in infants showed identity to different species of *Escherichia* phages (22.3% of abundance), *Lactococcus* phages (10.9%), *Salmonella* phage (8.2%), *Streptococcus* phages (5.8%), *Klebsiella* phages (5.3%), *Staphylococcus* phages (1.8%), *Clostridium* phages (3.4%) and *Enterococcus* phages (4.4%), with more than 50% of prevalence (Fig 6). Of note, *Salmonella* phage IME207, which was identified abundantly, is a novel lytic species isolated from *Klebsiella pneumoniae* which can lyse both *Klebsiella pneumoniae* and *Salmonella* [38]. On the other hand, in mothers, another crAssphage species, classified as uncultured, was identified as the second most abundant (10.3%), followed by Gokushovirus WZ-2015a (8.4%), *Actinomyces* virus Av1(4.1%), *Bacillus* virus G (2.3%) and human gut *Microviridae* SH-CHD12 (1.5%). Regarding the bacterial phage hosts, *Escherichia* (22.1%), *Bacteroidetes* (13.9%), *Lactococcus* (10.2%), *Klebsiella* (6.9%), *Streptococcus* (5.7%) and *Enterococcus* (4.3%) were the most commonly found, followed by *Clostridium*, *Enterobacter*, *Serratia* and *Staphylococcus*, each representing less than 3% of the bacterial reads found (S3 Fig).

### Alpha and beta diversity

Diversity estimations were performed using the normalized matrixes of viral read counts at the species level, and their association with factors of metadata (Fig 7). In infants, the Shannon alpha diversity of eukaryotic viruses was 1.4 ± 0.7 (Fig 7A) and of bacteriophages 3.2 ± 0.8 (Fig 7B), while in mothers these values were 0.8 ± 0.7 and 1.6 ± 0.8, respectively. In the diversity analysis, we found: i) the second semester of life of the infants was significantly richer (p=0.01) and more abundant (p=0.03) in eukaryotic viruses than the first semester (Fig 7A); ii) in contrast to this, phages were more abundant in the first semester (Fig 7B, p=0.02), iii) the diversity of bacteriophages was greater in infant 5 when compared to the other two infants, while in the eukaryotic virome only the richness was increased in this infant, iv) bacteriophages were significantly richer (p = 0.02) and more diverse (p = 0.05) in infants as compared to mothers (Fig 7B), as has been found in previous studies [5,7,29,31–33]; v) the eukaryotic virome, contrary to the phageome, was more abundant in mothers than in infants (Fig 7A, p=0.05).

**Fig 7.**
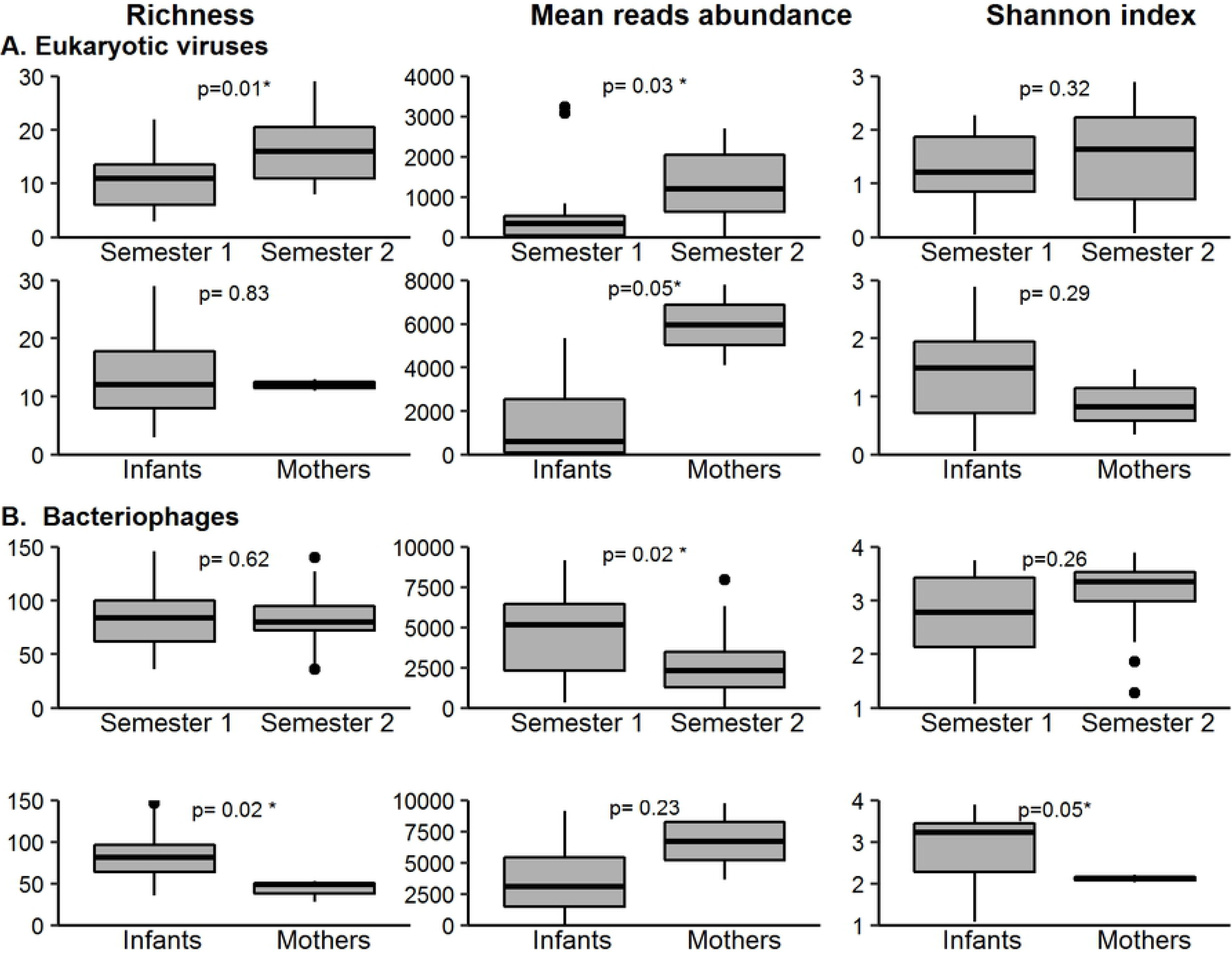
Bar plots representing viral diversity metrics as Chao richness index, mean abundance per species and Shannon diversity index. The metrics of the first vs. the second semester of infants, as well as the metrics of mothers as compared to those of infants, are compared. A) Eukaryotic viral diversity metrics. B) Bacteriophage diversity metrics.

The beta diversity of the eukaryotic viruses and of bacteriophages was calculated among mothers and infants, to individually compare them and to discern if there were patterns in association with metadata. Fig 8 illustrates the result of a Multidimensional Scaling (MDS) analysis using this beta diversity. Statistical analysis showed agreement with alpha diversity, obtaining significant difference between phage communities of the samples from mothers and infants (PERMANOVA using Adonis p = 0.002); however, such difference was not seen among eukaryotic viruses (p = 0.1). In both types of viruses, there was a difference between infant 5, who is a girl, and the other two infants (2 and 4) (p = 0.003), who are boys. Regarding age, there was a difference between eukaryotic viruses in the first and second semester of life (p = 0.01), but such difference was not observed when the phages were analyzed (p = 0.09).

**Fig 8.**
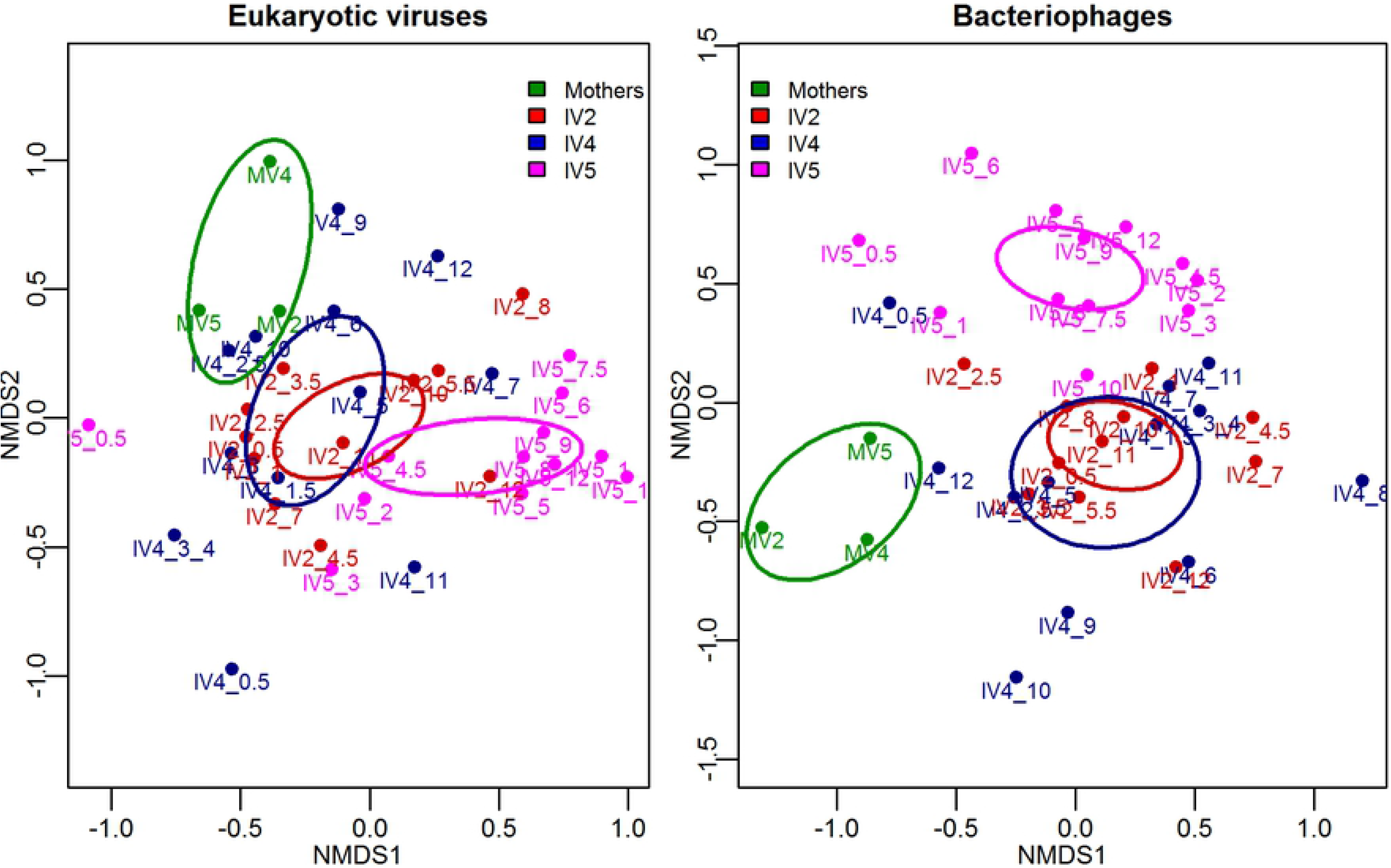
Multidimensional Scaling (MDS) analysis of eukaryotic viruses and bacteriophages at the species level. Bray-Curtis dissimilarity distances from normalized counts were used. Each point corresponds to a sample, and ellipses represent the standard errors of the centroids of the types of samples (mothers, infant 2, infant 4 and infant 5). Ellipses were calculated using the Ordiellipse function of the R package ‘vegan’ [24], at 95% confidence.

### Bacteria taxonomy composition

Even though the samples were filtered through 0.45 filters before nucleic acid extraction, to reduce bacterial contamination, about 49.4% of the total reads were classified as bacterial. In infants, the main phyla represented in the samples were Firmicutes (on average, 41.4% of the bacterial reads) and Proteobacteria (34.6%), followed by Bacteroidetes (14.6%) and Actinobacteria (8.3%) (S1 Fig). The remaining 31 phyla identified in infants represented only 1.1 % of the reads. In mothers, Firmicutes was also the most abundant phylum, with 55.9% of the reads, but the proportion of Bacteroidetes (36.6%) increased, while Proteobacteria (4.5%) and Actinobactaeria (1.3%) decreased as compared to infants. At the family level (S2 Fig), the most abundant and frequently identified in infants were *Enterobacteriaceae* (23.2%) and *Pseudomonadaceae* (5.6%) from Proteobacteria phylum; *Enterococcaceae* (10.0%), *Veillonellaceae* (5.9%), *Streptococcaceae* (4.2%), *Peptostreptococcaceae* (3.3%), *Clostridiaceae* (2.6%), *Lachnosipiraceae* (2.6%), *Lactobacillaceae* (2.4%) and *Staphylococcaceae* (2.0%) from Firmicutes phylum and *Bifidobacteriaceae* (7.5%) and *Prevotellaceae* (2.6%) from Actinobacteria and Bacteroidetes phyla, respectively. On the other hand, in mothers, the most abundant bacterial families were *Ruminococcaceae* (19.3%), *Clostridiaceae* (6.5%), *Lachnospiraceae* (5.6%), *Erysipelotrichaceae* (4.5%), *Peptostreptococcaceae* (2.6%) and *Veillonellaceae* (2.0%) from Firmicutes phylum; *Prevotellaceae* (11.7%), *Tannerellaceae* (8.1%), *Bacteroidaceae* (7.4%) and *Dysgonamonadaceae* (3.4%) from Bacteroidetes phylum and *Chlamydiaceae* (16.2%) from Chlamydiae phylum. Not surprisingly, the bacterial hosts of the most abundant and persistent phages in infants, described in the bacteriophages section, were identified (S3 Fig), followed two relation patterns in their abundance: a) Positive correlated such as Escherichia (15.9% on average per sample), Enterococcus (13.9%), Klebsellia (6.1%), Staphylococcus (4.9%) and Clustridium (1.8%) which corresponding phages had similar abundances. b) Negative Correlated such as Bacteroidetes (5.9%), and Lactoccocus (0.6), which phages are more abundant.

## DISCUSSION

It is of upmost importance to understand the way the enteric virome develops during infancy and what impact on the development of the gastrointestinal tract and the human health it may have. Few studies have described the gut virome of infants during the first year of life [5–7,29,31–33,39,40], with even fewer studies carried out in healthy children, in the community [5–7,29]; only two of these studies have been longitudinal. In this work, we characterized the monthly gastrointestinal virome, prokaryotic and eukaryotic, of three healthy infants during their first year of life.

Regarding eukaryotic viruses, 33 families were identified in infants, but only nine were frequently found and made up to 97% of all eukaryotic viral reads identified. Aside for plant-infecting viruses in the *Virgaviridae* family, the most abundant and frequently found were viruses belonging to the families *Anelloviridae*, *Astroviridae*, *Caliciviridae*, *Genomoviridae*, *Parvoviridae, Picornaviridae* and *Reoviridae*. Some viruses in these families are common human pathogens, especially in children; it is thus surprising that they were commonly found in the absence of gastrointestinal symptoms all along the year of the study. The gut mucosa of infants is under a process of maturation and receptors to pathogens may still be absent; although other immunological or nutritional factors may also be involved.

In our study, members of the *Anelloviridae* family were highly abundant, containing different species of torque teno mini virus and unclassified species in up to 80% of the samples. In previous studies, viruses from this family have been identified as the most abundant and frequent in healthy children [6,29,31,33,41–45], being more abundant during the first year of life [5,29,42], after which the abundances decrease. Their presence has been associated with a reduced host immune status; a higher abundance have been reported in patient with lung transplantation [46,47], AIDs [48], pulmonary diseases [49,50], cancer [51], among others; although their role in the pathogenesis of these diseases remains unclear [52]. Our results showed that anelloviruses were significantly more abundant in the second semester of life compared with the first one (P-value 0.01, S6 Table), specially TTV like mini virus and torque teno virus species. These results agree with Lim et. al. [5] and suggest that infants come more in contact with these viruses a few months after birth, from an unknown source. Viruses in the *Picornaviridae* family have also been frequently found in healthy children [9,41,43,44]. In our study, this family was identified in 80% of the infant samples, with parechovirus A and enterovirus A being the most common species. The duration of parechovirus secretion in the stool of healthy infants has been reported to last between 41 and 93 days [53]; in this regard, we also detected parechoviruses in infant feces during two consecutive months, followed by periods of null or undetectable levels.

Viruses in the *Caliciviridae* family have been found at a low frequency (7%) in healthy infants in metropolitan areas of the USA [5], while they were absent in an urban city but frequent (45%-60%) in rural communities of Venezuela [43] and Ethiopia [41]. In case-control studies they have been more frequently identified in sick as compared to healthy children [32,45,54]. In this context, it is remarkable that these viruses were present in up to 72% of the infant samples we studied and in the absence of gastrointestinal symptoms. Like anelloviruses, caliciviruses were significantly more abundant in the second semester as compared to the first six months of life. The virus species most identified were Norwalk and Sapporo viruses, found in 70% and 20% of the samples, respectively, and they showed a high level of genetic diversity. We were able to assemble seven complete Norwalk virus genomes and they belonged to genotypes GI and GII; we also assembled two Sapporo virus genomes which belonged to the GII genotype. A more detailed description of the genetic variability of these viruses will be described elsewhere (Rivera-Gutiérrez et al., in preparation).

We identified a large set of 27 species of *Parvoviridae* and *Genomoviridae*, with most of them being insect and animal viruses. They were sporadically identified and in low abundance, possibly reflecting environmental contamination, except for human bocaviruses, which were identified in 11 of 35 infant samples. Such high prevalence was not surprising, as bocaviruses have been previously reported in feces of more than 40% of asymptomatic children [55]. Rotavirus A, a common etiological agent of infantile gastroenteritis, belonging to the *Reoviridae* family, was identified in 66% of the infant samples. Interestingly, all rotavirus reads detected showed an identity of 100% with different genes of the RotaTeq vaccine strains. The rotavirus vaccine was administered to the three infants at around two, forth, and six months of age, except for infant 5 who did not receive the last dose. Surprisingly, we identified the rotavirus vaccine strain in 4 out of 5 samples just before their first vaccination, which suggests a frequent transmission of the vaccine strain by close contact with vaccinated people, as it has been suggested in a previous study of transmission from vaccinated infants to their unvaccinated co-twin [56]. Rotavirus A was more abundant in the second semester (p-value 0.001, S6 Table), when the three doses had already been administered to infants.

Regarding plant-infecting viruses, those in the *Virgaviridae* family were frequently detected in the three children along the year of study. The *Tobamovirus* genus was the most frequent, with tropical soda apple mosaic virus, pepper mild mottle virus, and opuntia tobamovirus 2 being the most common species. Our results showed a large diversity of tobamoviruses circulating in the population, suggesting that infants are continuously exposed to an extensive and dynamic collection of these plant viruses, even before infants begin to ingest food other than mother’s breastmilk, including baby formula or other liquids, indicating a distinct source of origin for these viruses. We recently reported the genetic diversity and dynamics of tobamovirus infection in infants, as wells as the potential implications of these findings [30].

The richness of eukaryotic virus species found in this work was higher than in previous studies in healthy children, in which less than 9 different virus species were found per individual samples [5,7,41,43,45]. We identified on average 15 (±9 s.d.) virus species per sample, which is higher even when compared to previous reports in rural or small village communities [41,43,45]. In general, we also identified a greater number of enteric viruses compared to previous studies carried out not only in healthy infants, but also in sick children [32,45]. Several factors may influence these results and should be considered in future studies. These include, an unbiased nucleic acid extraction method, which does not target only DNA viruses; depth of the sequencing carried out; socio-economic or demographic characteristics of the community or even a greater susceptibility or exposure of infants in our community as compared to other populations. When virus diversity in our samples was analyzed, it was found that the eukaryotic virome significantly increased in richness and abundance during the second semester of life, suggesting eukaryotic viruses are established as result of an increased environmental exposure of infants with age, in agreement with previous observations [5]. In line with this observation, viruses in the *Anelloviridae*, *Caliciviridae*, *Reoviridae* and *Virgaviridae* families were found more abundantly in the second semester, as compared to the first six months of life. It is important to mention that we estimated diversity based on annotated taxa, not at contig level, since in our experimental procedure virus-like particles were not purified; and thus, our values cannot be compared with those of previous studies that use this method.

Most viral reads in this work were assigned to bacteriophages, both in infants (84.5 ∓ 24%) and mothers (61.4 ∓ 37%). Unlike eukaryotic viruses, and in contrast to previous findings [5], no difference in richness was found between the first and second semester of life, although the mean abundance was greater in the first semester. Of note, bacteriophages were significantly richer and more diverse in infants than in their mothers, which agrees with a previous study where the richness and diversity of bacteriophages in the infant gut virome are reported to be higher than in adults, and decreases with age [44]. The dominant phages belonged to the *Siphoviridae*, *Myoviridae* and *Podoviridae* families in the *Caudovirales* order. Although previous studies have reported that the majority of gut bacteriophages seem to engage in lysogenic interactions with their hosts [35,57–59], in our study the particular dominant phages in all samples were the recently described CrAssphages, an expansive diverse group of lytic bacteriophages with podovirus-like morphology that includes the most abundant viruses from the human gut [60]. In this regard, it is important to point out that these viruses have been reported to stably infect bacteria within the phylum Bacteroidetes during long periods of time both, *in vitro* and *in vivo* [37], although the mechanisms underlying this unusual relationship of carrier state-type are unknown. In any case, this type of interaction may start early in life, at least in the studied community.

CrAssphages have been reported to represent up to 95% of the total viral load in the adult’s gut, and to be present in 73% to 77% of samples analyzed in diverse human populations [60,61]. Recent studies have shown that these viruses can be found as early as one week after birth [62] and it has been suggested that they could be vertically transmitted from mother to child [63]. In our study these phages were detected in 86% of the infants samples, with abundances ranging between 1% and 82% and with up to 96% of abundance in mothers; this frequency of detection was higher than that found in previous studies in infants, where they were detected in up to 53% of the samples [61]. Although this group of phages are diverse, the crAssphages identified in this work had homology only to crAssphage Azobacteroides phage, Bacteroides phage, Cellulophaga phage, IAS virus and one uncultured crAssphage. Interestingly, at 15 days of age, the crAssphages were more abundant (on average 35%) than the Bacteroidetes (12%), a trend that was maintained during the first semester and in a ratio that started to change during the second semester of life, becoming the inverse, and by month 10 the Bacteroidetes phages were more abundant than the CrAssphages (S4-A Fig). More data is needed to understand the roles and dynamics of CrAssphage-bacteria in gut equilibrium. Other virulent *E*. coli phages in the *Myoviridae* and *Podoviridae* families were abundantly identified in all samples, persisting for prolonged periods of time (S4-B Fig). These phages infect *Escherichia* spp and *Klebsiella* spp. Also, apart from abundantly identifying proteins associated with lysogenic viruses such as integrases, we identified portal proteins which are used by lytic phages to form a pore that enables DNA passage during their packaging and ejection. These results suggest that there is a core of virulent bacteriophages in early life of humans as there is in healthy adults [64].

The results and conclusions of this study are limited by a small sample size, although our primary goal was to have a first glimpse on the composition and dynamics of the eukaryotic and prokaryotic virome of healthy Mexican infants in a rural community during first year of life. Our findings suggest the existence of an early and constant contact of infants with a diverse array of eukaryotic viruses, whose composition changes over time. In addition to the phageome, that seems to be well established given the ubiquitous presence of their microbial hosts, the eukaryotic virus array could represent a metastable virome, whose potential influence on the development of the infant’s immune system or on the health of the infants during childhood, remains to be investigated.

## Acknowledgments

This work was partially supported by grants 272601 and 257091 from the National Council for Science and Technology-Mexico (CONACYT), and by grants IG200317 and IN215018 from DGAPA-PAPIIT/UNAM. We are grateful to Enrique Gonzales, Eric Hernández, Miriam Nieves, and Marco A. Espinoza for their excellent technical assistance. A special acknowledgement for Xochiquetzalli Soto-Martínez for her invaluable work performed with the families in the community. We also thank Jerome Verleyen and Juan Manuel Hurtado for their computer support.

## Supporting information

**S1 Fig. Relative abundance of the sequence reads from the most frequent bacterial phyla across all samples**.

**S2 Fig. Relative abundance of the sequence reads from the most frequent bacterial families across all samples**.

**S3 Fig. Relative abundance of the sequence reads from the most frequent bacterial species across all samples**.

**S4 Fig. Average abundance of reads of the two most represented phages and their bacteria hosts in the infants, all year long**.

**S1 Table. Characteristics of mother-infant binomials**.

**S2 Table. Sampling summary and sequencing data generated for each sample**.

**S3 Table. Reads abundance of eukaryotic viral species**.

**S4 Table. Reads abundance of prokaryotic viral genera**.

**S5 Table. Reads abundance of prokaryotic viral species**.

**S6. Table. Taxonomic differential abundance between groups at different levels**

